# Discovery of Type IV filament membrane alignment complex homologs in *H. pylori* that promote soft-agar migration

**DOI:** 10.1101/2023.04.27.537399

**Authors:** Jashwin Sagoo, Samar Abedrabbo, Xiaolin Liu, Karen M. Ottemann

## Abstract

The stomach pathogen *Helicobacter pylori* utilizes two scaffold proteins, CheW and CheV1, to build critical chemotaxis arrays. Chemotaxis helps bacteria establish and maintain infection. Mutants lacking either of these chemotaxis proteins have different soft agar phenotypes: deletion of *cheW* creates non-chemotactic strains, while deletion of *cheV1* results in 50% loss of chemotaxis. In this work, we characterized the *cheV1* deletion mutant phenotype in detail. *cheV1* deletion mutants had poor soft-agar migration initially, but regained migration ability over time. This improved bacterial migration was stable, suggesting a genetic suppressor phenotype, termed Che+. Whole-genome sequencing analysis of four distinct *cheV1* Che+ strains revealed single nucleotide polymorphisms (SNPs) in a common gene, HPG27_252 (HP0273). These SNPs were predicted to truncate the encoded protein. To confirm the role of HPG27_252 in the *cheV1* phenotype, we created a targeted deletion of HPG27_252 and found that loss of HPG27_252 enhanced soft-agar migration. HPG27_252 and CheV1 appear to interact directly, based on bacterial two-hybrid analysis. HPG27_252 is predicted to encode a 179 amino acid, 21 kDa protein annotated as a hypothetical protein. Computational analysis revealed this protein to be a remote homolog of the PilO Type IV filament membrane alignment complex protein. Although *H. pylori* is not known to possess Type IV filaments, our analysis showed it retains an operon of genes for homologs of PilO, PilN, and PilM, but does not possess other Type IV pili genes. Our data suggest the PilO homolog plays a role in regulating *H. pylori* chemotaxis and motility, suggesting new ideas about evolutionary steps for controlling migration through semi-solid media.

## Introduction

Bacteria use motility to succeed in complex environments and infections, responding to both harmful and helpful conditions and ultimately moving towards beneficial conditions (Wuichet & Zhulin, 2010). Chemotaxis advances bacterial colonization during infection and aids in their survival in various environments (Ottemann & Miller, 1997; Szurmant & Ordal, 2004; Wadhams & Armitage, 2004). Chemotaxis signal transduction is necessary for this bacterial response. In the basic system, bacteria sense conditions using chemoreceptors, and relay this information to the flagellar motor via a set of signal transduction proteins. The basic set of chemotaxis signaling proteins is conserved throughout bacteria, but there are variations in terms of accessory proteins that decorate each system. The study of these accessory proteins and non-canonical systems promises to provide insights into how chemotaxis signaling systems work (Adebali *et al*., 2017; Kennedy *et al*., 2022; Liu & Ottemann, 2022; Russell *et al*., 2013).

Chemotaxis signaling occurs when chemoreceptors sense environmental parameters and control an associated kinase. The chemoreceptors and kinase do not directly associate but instead use scaffold proteins, also called coupling proteins, to make the connection (Alexander *et al*., 2010; Vass *et al*., 2022). The chemotaxis scaffolds are the proteins CheW or CheV1. CheW proteins are small, two-domain proteins that carry out protein-protein interactions. One type interacts with both chemoreceptors and CheA, to link them, and the other type occurs between CheW monomer to connect them. CheV proteins, in comparison, have CheW domains fused to phosphorylatable response regulator domains, but are less understood (Alexander *et al*., 2010; Vass *et al*., 2022). CheV proteins, however, have been shown to be capable of promoting both types of interactions, and additionally may play a role as a dead-end phosphorylation site, a so-called phosphate sink (Abedrabbo *et al*., 2017; Alexander *et al*., 2010). However, there are gaps in our understanding of CheV function as compared to CheW.

One bacterium that relies on chemotaxis signaling during its lifestyle is the pathogen *Helicobacter pylori*. This microaerophilic Gram negative microbe colonizes the human stomach, leading to gastritis, peptic ulcer disease, and stomach cancer in some individuals (Crowe, 2019). The ability of *H. pylori* to colonize the stomach is modulated by motility and chemotaxis, in addition to multiple other virulence and colonization factors. *H. pylori* uses chemotaxis to achieve rapid stomach colonization and to interact with the host in a way that modulates inflammation (Johnson and Ottemann, 2017). Gaining an understanding of *H. pylori* chemotaxis provides insights into bacterial-host interactions and pathogenesis (Fung *et al*., 2019; Keilberg *et al*., 2016; Rolig *et al*., 2011; Terry *et al*., 2005; Williams *et al*., 2007).

*H. pylori* has a relatively large number of scaffold proteins, including one CheW and three CheV proteins that all promote normal chemotaxis to varying degrees (Alexander *et al*., 2010; Lowenthal *et al*., 2009; Pittman *et al*., 2001). *cheW* mutants have severe soft agar migration defects and a strong smooth-swimming bias (Lowenthal *et al*., 2009; Pittman *et al*., 2001). CheW is essential for forming chemotaxis protein complexes (Abedrabbo *et al*., 2017). *cheV1* mutants similarly have a similar strong smooth swimming bias and CheV1 is also essential for forming chemotaxis signaling complexes (Abedrabbo *et al*., 2017; Lowenthal *et al*., 2009). *cheV1* mutants, in contrast, have only a ∼50% decrease in soft agar migration (Lowenthal *et al*., 2009; Pittman *et al*., 2001). We were thus struck by this apparent discrepancy. We noticed, however, that the previous chemotaxis soft agar assays were all done with relatively long incubation times, when genetic chemotaxis-able suppressors have been documented to arise (Terry *et al*., 2006). We therefore wondered whether the chemotaxis phenotype of *cheV1* mutants might have been mischaracterized. We reassessed the phenotypes of *cheV1* mutants and found that these mutants behave on soft agar in a manner that is consistent with full loss of chemotaxis but with ready genetic suppression. Che^+^ suppressor arose during incubation and possessed near wild-type levels of soft agar migration. Whole genome sequencing identified that these Che^+^ suppressors had mutations in an uncharacterized open reading frame HPG27_252, a gene which was annotated as a hypothetical protein with unknown function. The mutations in this gene caused either a truncation or a frame shift which resulted in the protein not being made correctly. The role of HPG27_252 was confirmed by targeted mutation, showing that elimination of this gene allowed recovery of soft agar migration. Computational analysis shows that HPG27_252 encodes a remote ortholog of PilO, part of the Type IV filament cytoplasmic alignment complex. Our work thus suggests that HPG27_252 encodes a previously unappreciated regulator of soft-agar motility.

## Results

### cheV1 mutants have initial soft-agar migration defects that lessen over time

CheV1 plays a pivotal role in forming chemotaxis arrays (Abedrabbo *et al*., 2017). Given this role, it was surprising that *H. pylori cheV1* null mutants have only a partial soft-agar migration defect (Lowenthal *et al*., 2009; Pittman *et al*., 2001). The *cheV1* null mutant soft-agar phenotype was therefore carefully evaluated using mutants in which nearly all of the *cheV1* open reading frame was deleted and replaced by a chloramphenicol resistance cassette (*cat*), referred to as Δ*cheV1*. These mutants were evaluated using the well-characterized soft-agar assay, in which in low percentage agar (<0.5% agar) allows motile, chemotactic bacteria to form expanded colonies. These expanded colonies have a large migration diameter compared to non-motile or non-chemotactic cells (Che^−^), which stay near the point of inoculation (Wolfe & Berg, 1989). The *H. pylori* Δ*cheV1* strain soft-agar colony diameter was thus measured over multiple days. After incubating four days, the Δ*cheV1* mutant had a substantial soft-agar defect that was significantly below WT and not different from a fully Che^−^ strain (Fig. 1A-B). The Δ*cheV1* mutant phenotype changed over time, however, with an increased colony diameter that became significantly different from the Che^−^ strains by day 7 (Fig. 1C-D). This outcome suggested that Δ*cheV1* mutants recovered some ability to migrate in soft agar. To assess whether this phenotype was genetically stable, we collected and purified single colonies from the outer edges of these expanded Δ*cheV1* soft agar colonies and screened these strains on separate soft-agar assays. The colony migration diameters of these colonies were variable but displayed a significantly larger migration diameter than the Δ*cheV1* parent strain or a fully non-chemotactic strain lacking all chemoreceptors (Δ*tlpABCD*) (Fig. 2). These results suggested that the Δ*cheV1* mutants may have acquired suppressor mutations that allowed for a significantly greater soft-agar migration compared to the parent Δ*cheV1* phenotype. These strains were termed Δ*cheV1* Che^+^ to indicate their recovered soft-agar migration phenotype.

**Fig. 1.**
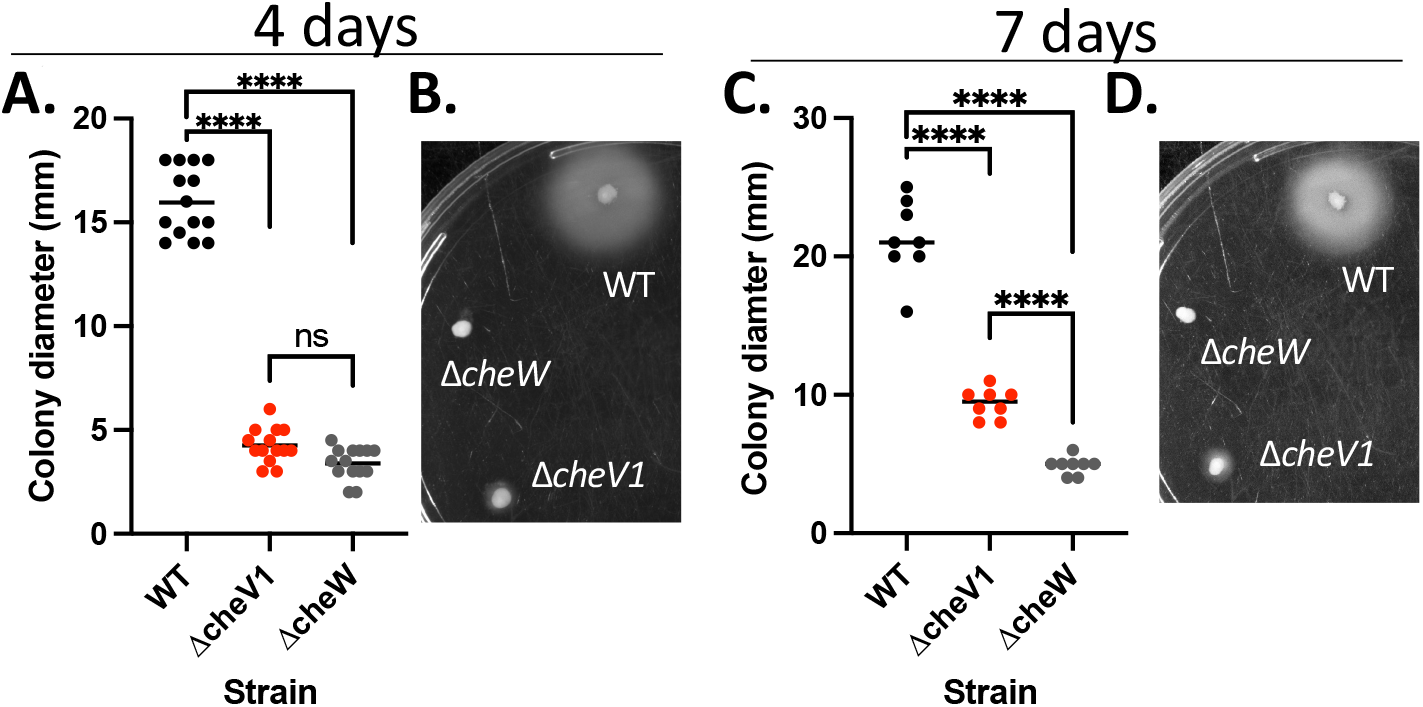
c*heV1* mutants have severe initial soft agar defects that lessen over time. Chemotaxis abilities were determined using a soft agar assay of wild-type G27 and its isogenic mutant strains. Bacteria were inoculated from blood plates with a pipette tip into Brucella broth-FBS 0.3% agar plates and incubated at 37°C under microaerobic conditions. (A-C) Quantification of colony diameters in soft agar plates after 4 or 7 days, as indicated. N = 8-14 plates/strain. Each point represents one measurement, and the bar indicates the mean. (B-D) Soft agar plate images of indicated *H. pylori* G27 strains after 4 or 7 days of incubation. Statistical analysis was done using one-way Anova. **** indicates p<0.0001.

**Fig. 2.**
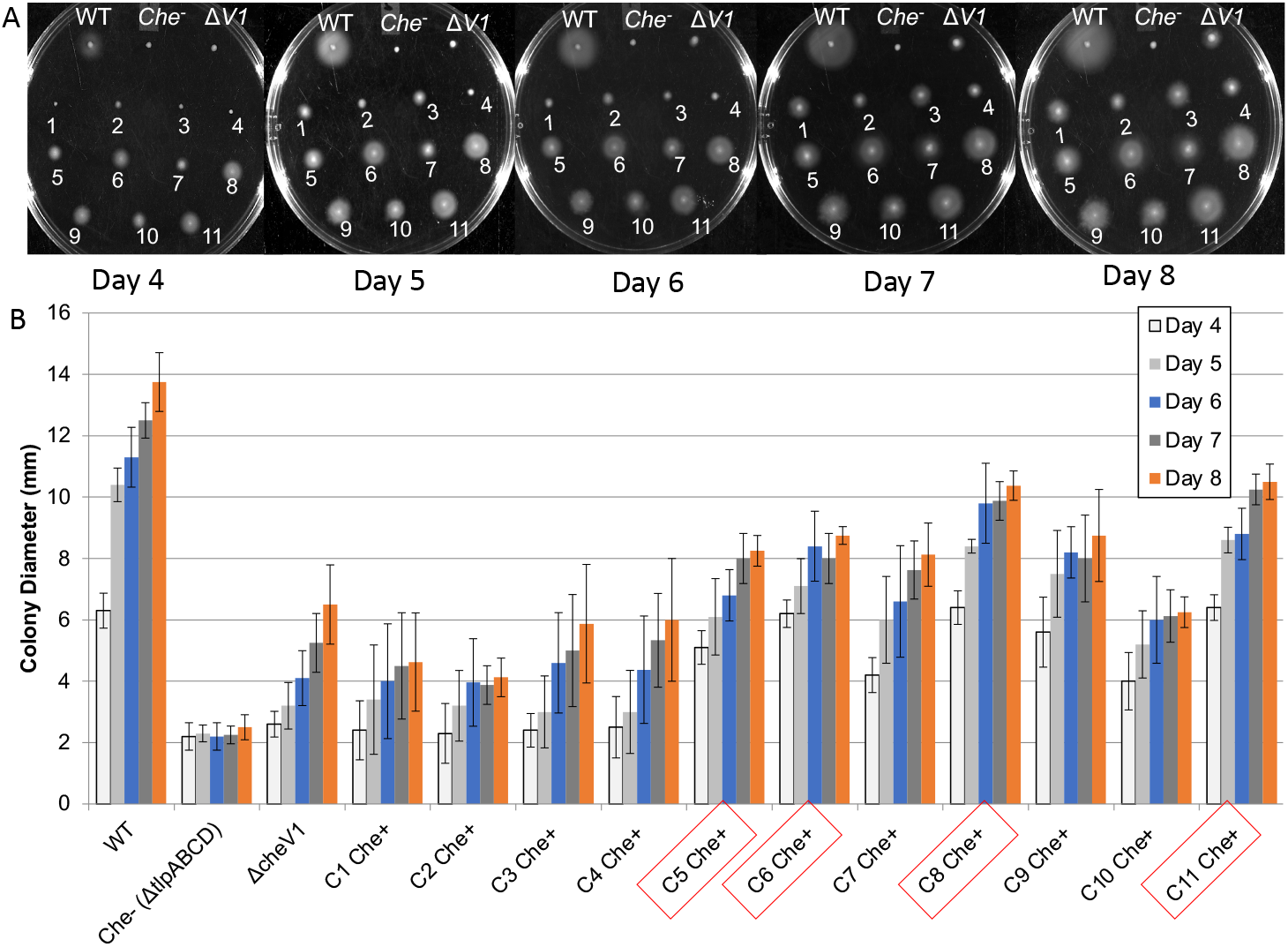
Chemotaxis assay of *cheV1* Che+ suppressor colonies. Chemotaxis abilities were determined in soft agar as described in Fig. 1. *cheV1* mutants that gained the ability to migrate in soft agar were collected from the outer edge of the bacterial growth halo of *cheV1* mutants grown on soft agar plates for 5-6 days. The bacteria on the pipette tip were streaked on blood plates to isolate single colonies, referred to as C1-11 Che+, and were retested. (A) Images of Brucella broth-FBS soft agar plates incubated for 4-8 days. Numbers indicate the Δ*cheV1* suppressor colonies. (B) Quantification of colony diameters in soft agar plates. Red boxes represent colonies sent for genome sequencing—C5, C6, C8, and C11. Data are shown as a mean and error bars represent standard deviation (n= 9-12 for all strains).

### cheV1 Che^+^ strains display normal growth and do not change reversal frequency in liquid

Recovery of soft-agar migration could arise through several mechanisms, as the migration relies on a combination of growth, nutrient utilization, motility, and chemotaxis. We thus evaluated whether growth was affected in the Δ*cheV1* Che^+^ mutants, and found it to be unaffected and similar to wild type and the Δ*cheV1* parent (Supplemental Fig. 1). We then examined chemotactic output, which can be evaluated by examining swimming reversals (Lertsethtakarn *et al*., 2015; Terry *et al*., 2006). Δ*cheV1* mutants displayed extremely few flagellar reversals compared to WT, when evaluated in liquid media (Supplemental Fig. 2), as reported previously in line with highly CCW-biased flagellar rotation (Antani *et al*., 2021; Lowenthal *et al*., 2009). Δ*cheV1* Che^+^ variants similarly had few reversals in liquid media, nearly the same low-reversal phenotype as the Δ*cheV1* mutant parent (Supplemental Fig. 2). Overall, these results suggested that the Δ*cheV1* Che^+^ suppressors did not acquire their soft-agar migration phenotype by affecting growth rates or flagellar reversals.

### Genomic sequencing of four Che+ suppressors reveal acquired SNPs in HPG27_252

Because the Δ*cheV1* Che^+^ phenotype was stable and thus consistent with a genetic change, we determined the genomic sequences of four Che^+^ suppressors (numbers 5, 6, 8 and 11 in Fig. 2) and the Δ*cheV1* parent. These sequences were compared to the published *H. pylori* G27 reference genome (Baltrus *et al*., 2008). Multiple single nucleotide (SNPs) were discovered across these suppressors when mapping the sequencing data to the reference genome (Supplemental Table 1) but one locus stood out because it had distinct SNPs in each of the four *cheV1* Che^+^ strains that were not present in the *cheV1* parent (Supplemental Table 1). These SNPs occurred in different positions in this locus resulting in truncated proteins (Fig. 3). These SNPs were confirmed by Sanger sequencing of PCR products of this region. This locus, HPG27_252 (HP0273 in the reference *H. pylori* 26695 genome), encodes a hypothetical protein that is conserved across *H. pylori* (Fig. 4A). Taken together, this data suggested that HPG27_252 is the site of mutations in Δ*cheV1* Che^+^ suppressors.

**Fig. 3.**
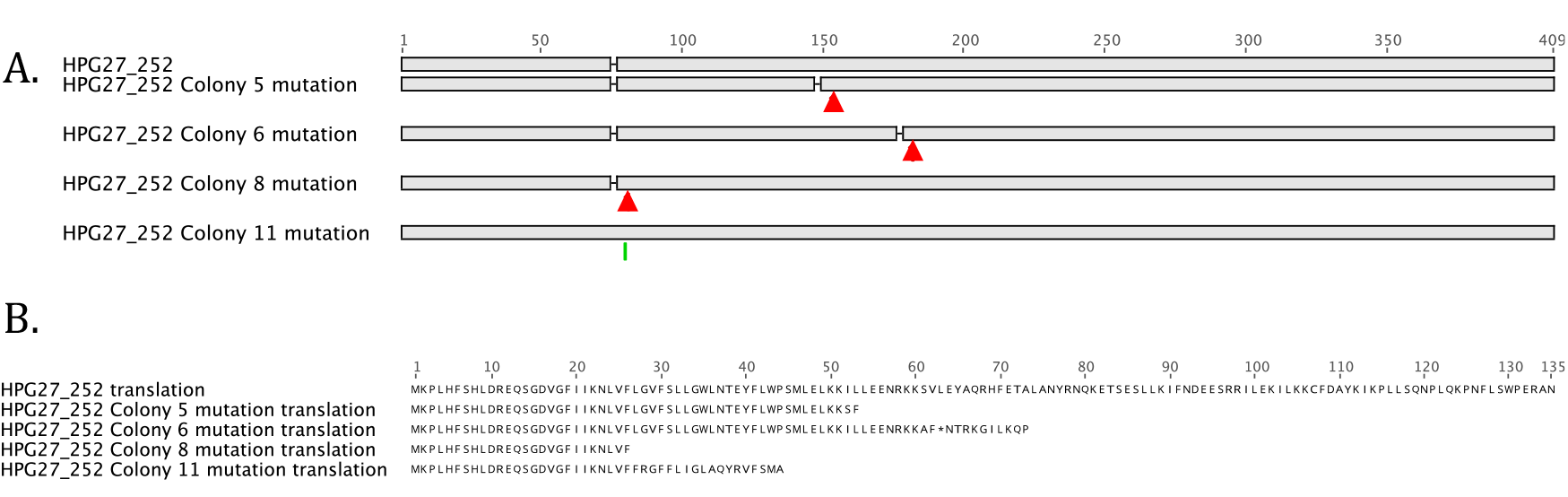
HPG27_252 is mutated in *cheV1* Che^+^ suppressor mutants. A. locations mutations in the HPG27_252 gene for colony 5 (deletion), 6 (deletion), 8 (deletion) and 11 (insertion). B. Representation of the translation of HPG27_252 WT, resulting a 135 amino acid protein, compared with the four mutations that all cause frameshifts that truncate the resulting product and in some cases alter the C terminal end. The corrected G27 HPG27_252 extends the C-terminal end with an additional 44 amino acids.

**Fig. 4.**
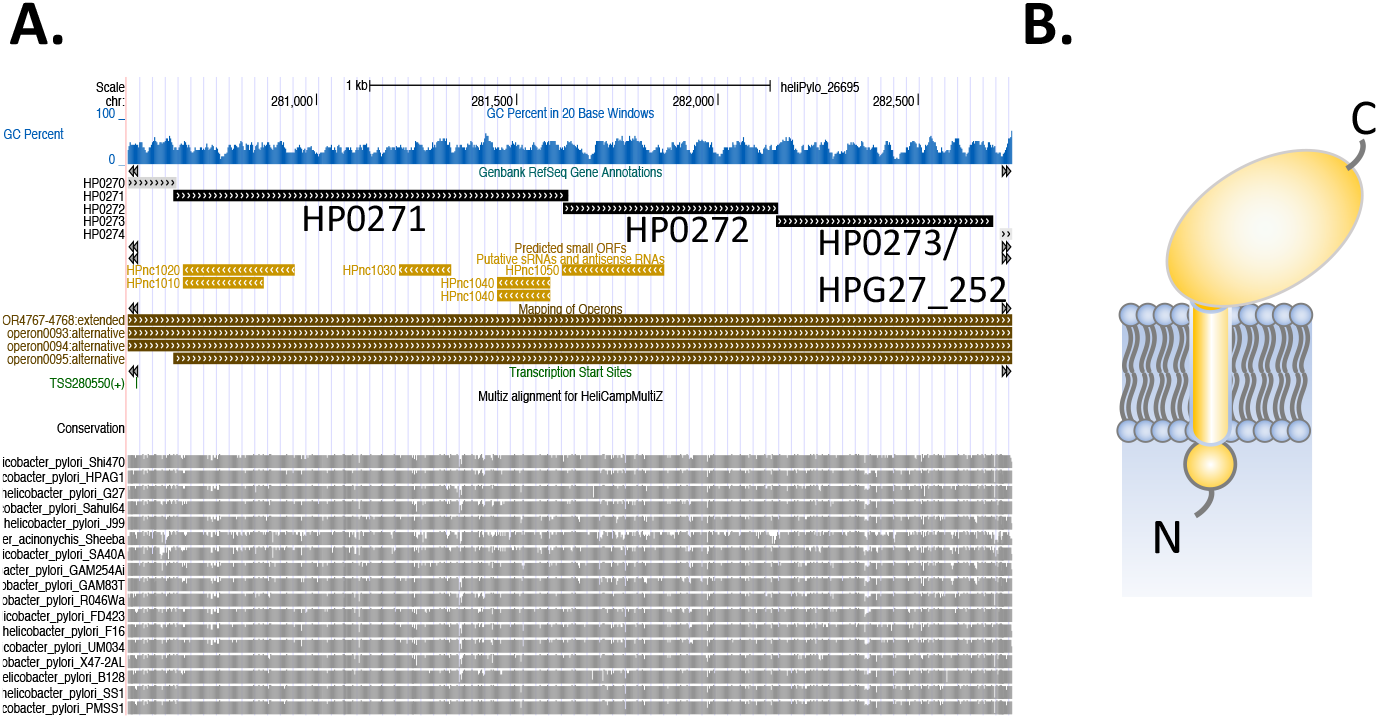
HPG27_252 (HP0273 in the *H. pylori* 26695 reference genome) is conserved across *H. pylori* and encodes a remote ortholog of PilO. A. Operon prediction in brown lines, based on transcriptome data from *H. pylori* 26695 (Sharma *et al*., 2010)shows that HP0273 is part of an operon with HP0272 (predicted PilN ortholog) and HP0271 (predicted PilM ortholog). Predicted small RNAs encoded in this region are shown in gold. At the bottom is conservation in other *H. pylori* and *H. acinonychis* strains, with the heights of the bars representing degree of conservation. (B). Predicted structure of HPG27_252 protein with an 24 amino acid N-terminal cytoplasmic domain, a transmembrane domain from amino acids 25-34, and a C-terminal periplasmic domain from amino acids 35-179.

### Elimination of HPG27_252 in the ΔcheV1 parent results soft agar migration recovery

Given that Δ*cheV1* Che^+^ suppressors have SNPs in HPG27_252 that truncated the protein, we explored whether loss of the HPG27_252 gene was sufficient to allow *cheV1* mutants to migrate in soft agar. We created an HPG27_252 loss of function allele by replacing all but the regions encoding the first 24 and last 19 basepairs of the coding sequence with a kanamycin resistance gene, and then creating strains that combined this mutation with the Δ*cheV1* one. The resulting strains were then inoculated in soft-agar plates. The swarm diameter of these strains was larger than the *cheV1* parent, confirming the role of this gene in the phenotype of the suppressors (Fig. 5).

**Fig. 5.**
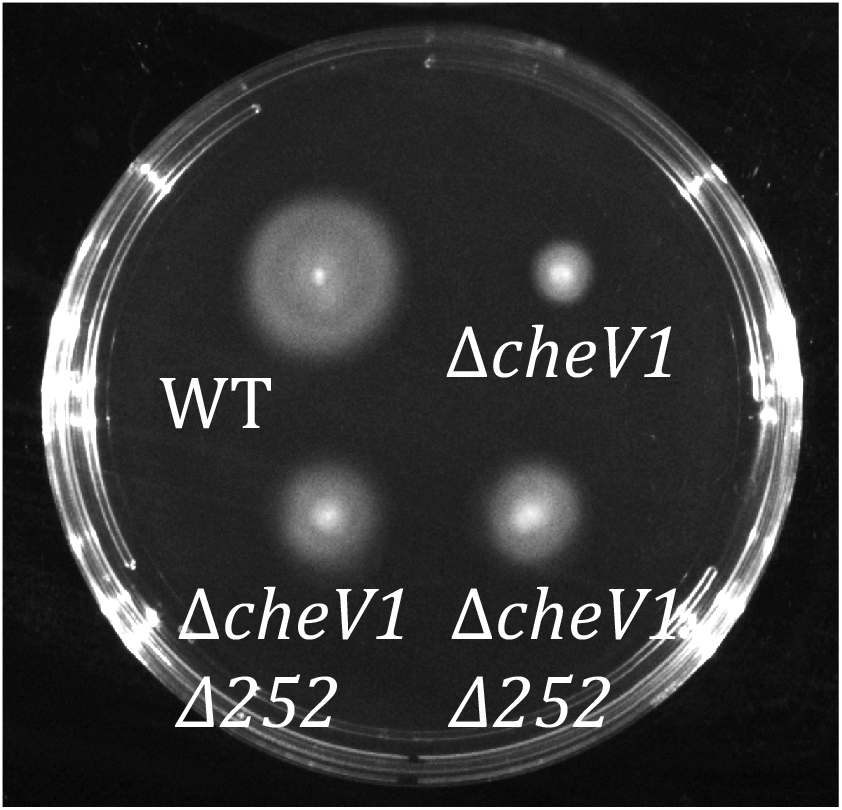
Loss of HPG27_252 in the Δ*cheV1* mutant allows soft agar migration recovery. HPG27_252 was deleted and replaced by a kanamycin resistance cassette in the *H. pylori* Δ*cheV1* background. Soft agar migration as assessed after 5 days, using Brucella Broth-FBS-soft agar plates. Duplicate Δ*cheV1* ΔHPG27_252 mutants (Δ*cheV1* Δ252) are shown. Plate is representative of over five assays.

### HPG27_252 encodes an inner membrane protein with remote homology to type four filament cytoplasmic membrane complex members

The HPG27_252 encoded protein was analyzed using bioinformatics tools, as it was annotated as a hypothetical protein in *H. pylori*. We noted that there is an error in the original G27 genome for this gene, of a missing adenine nucleotide, which truncated the protein. Data from both the *cheV1* mutant parent and the four suppressor colonies supported that there was an error in the reference genome. The corrected sequence revealed a longer open reading frame of 179 amino acids, similar to annotations for this gene in other *H. pylori* genomes. The encoded full length protein has a predicted 24 amino acid N-terminal region, a predicted transmembrane domain from amino acids 25-34, and a C-terminal periplasmic domain from amino acids 35-179 (Fig. 4B). BLAST and BLASTp yielded no significant hits to known function proteins, whereas PFAM identified only an uncharacterized DUF, DUF5401. We therefore utilized HHPred searches to seek remote homology based on statistical models and found remote homologs PilN and PilO (Table 1).”

**Table 1:** HPG27_252 homology to PilO and PIlN. Remote homologs of HPG27_252, HPG27_251, and HPG27_250 using HHPRED at the MPI Bioinformatics Toolkit (Zimmermann *et al*., 2018). Hit indicates the matched protein ID in the RCSB protein data bank (https://www.rcsb.org/structure/3JC8). Hit name indicates a short description of the matched protein. Probability and E value indicate the estimated probablity of that the template hit is homologous to the query. Probability scores are calculated to include the secondary structure similarity, with scores greater than 95% indicating the homology is highly certain. E value is an alternative measure of statistical significance that does not include the secondary structure, and is therefore described as less sensitive than the probability score. Aligned indicates the region in the query that matched the hit.

| Query | Hit | Hit Name | Prob | E-value | Aligned |
| --- | --- | --- | --- | --- | --- |
| HPG27_252/HP0273 | 3JC8_NI | PilN *Myxococcus xanthus* DK 1622 | 98.9 | 1.7E-06 | 47-177 |
|  | 3JC9_Oi | PilO *Myxococcus xanthus* DK 1622 | 98.9 | 6.7E-06 | 20-177 |
| HPG27_251/HP0272 | 3JC8_NI | PilN *Myxococcus xanthus* DK 1622 | 99.73 | 3.8e-15 | 15-139 |
|  | 4BHQ_B | PilN *Thermus Thermophilus* | 99.43 | 3.3e-11 | 44-146 |
|  | 3JC9_Oi | PilO; *Myxococcus xanthus* DK 1622 | 99.13 | 1.2e-8 | 25-52 |
| HPG27_250/HP0271 | 3WT0_C | FtsA; Cell division protein<br>*Staphylococcus aureus* | 92.53 | 11 | 1-305 |
|  | 3HRG_A | BT_3980 uncharacterized protein with<br>actin-like ATPase fold; *Bacteroides*<br>*thetaiotaomicron* BT_3980, | 91.93 | 12 | 19-191 |
|  | 2YCH_A | PILM *THERMUS THERMOPHILUS* | 90.57 | 17 | 1-305 |

PilO, PilN and related proteins like GspM are part of the cytoplasmic membrane complex of Type IV filament (TFF) nanomachines, typified by Type IV pili (T4P) and Type II secretion systems (T2SS) (Leighton *et al*., 2016; McCallum *et al*., 2019; Michel-Souzy *et al*., 2018). Both of these types of filaments form structures that extend from the cytoplasmic membrane to outside the cell. They serve various functions, including adhesion, extension-retraction based motility, and protein secretion. *H. pylori* is not known to have any T4P or T2SS. However, in support of the idea that HPG27_252 encodes a bona fide PilO homolog, we analyzed the gene neighborhood and found that the genes that are predicted to be co-transcribed with HPG27_252 also encode predicted components of the T4P cytoplasmic membrane complex, PilN and PilM, with an overlapping gene arrangement that suggests the protein products may interact (Fig. 4A). This result provides further evidence that HPG27_252 encodes a homolog of TFF proteins.

### Bacterial two-hybrid analysis supports a direct interaction between CheV1 and HPG27_252

Genetic suppressors often indicate interactions between its associated protein(s). We wanted to test this interaction hypothesis to see if HPG27_252 had an interaction with CheV1directly using the bacterial two hybrid assay (BACTH). For this analysis, full length HPG27_252 was cloned into the BACTH vectors, with the adenyl cyclase fragments at either the N-(pUT-252) or C-terminal end (pUTC-252). These plasmids were then used with previously constructed full length CheV1 vectors (Abedrabbo *et al*., 2017) and interactions evaluated on LB agar + Xgal. A strong interaction was detected between HPG27_252 and CheV1, but only when the adenyl cyclaseT18 fragment was at the N-terminal end (Fig. 6). This end is predicted to be cytoplasmic domain of HPG27_252, and thus the interactions supports this orientation. This finding suggests the two proteins interact via the small N-terminal cytoplasmic domain of HPG27_252, adding credence to the hypothesis that HPG27_252 was indeed responsible for the recovery of migration diameters of the four suppressors in the soft-agar assay.

**Fig. 6.**
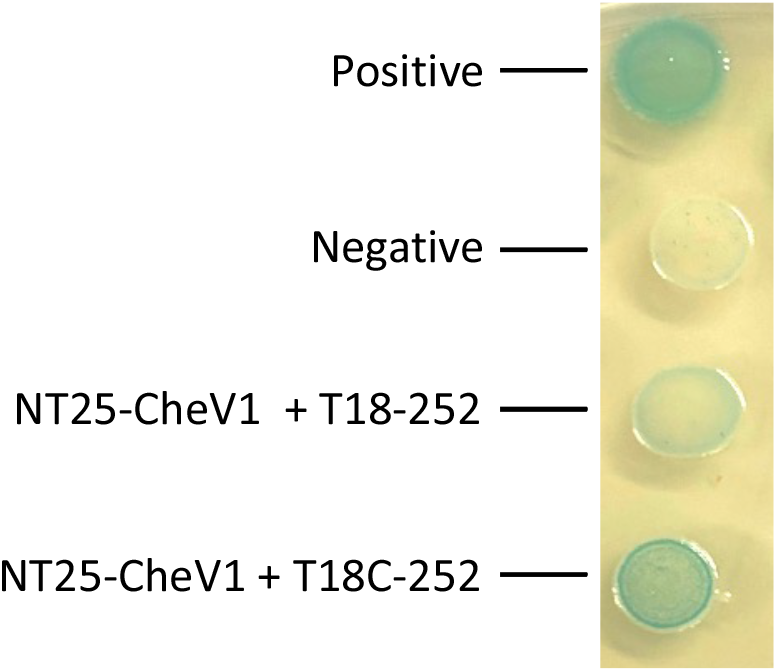
BACTH analysis of interactions between CheV1 and HPG27_252. *cheV1* was cloned into the pKNT25 (NT25) (Abedrabbo *et al*., 2017), and HPG27_252 was cloned into pUT18 (T18) or pUT18C (T18C). Data shown are representative of 3 replicates.

## DISCUSSION

We report here the identification of a new player in *H. pylori* soft agar migration—the product of an uncharacterized gene, HPG27_252. Mutants lacking this gene allow enhanced soft agar migration of *cheV1* mutants. HPG27_252 and its orthologs in other *H. pylori* genomes are annotated as hypothetical proteins. Our analysis here suggests this gene’s product, as well as the others in its operon, represent remote homologs of the Type IV filament cytoplasmic membrane alignment complex.

We report here that *cheV1* mutants have substantial soft agar migration defects, akin to fully non-chemotactic strains. The *cheV1* soft agar migration phenotype prior to this work was unclear. *cheV1* mutants had previously been reported to have ∼ 40% ability to migrate in soft agar (Lowenthal *et al*., 2009; Pittman *et al*., 2001). *H. pylori* soft agar assays, however, are incubated for 4-8 days, a time during which second site suppressor mutations have been reported to arise (Terry *et al*., 2006). Indeed, we noticed that *cheV1* mutants had a stronger early defect than later. This observation provided the impetus for this work, to carefully characterize the *cheV1* mutant phenotype. Indeed, we found that *cheV1* mutants are equivalent to other non-chemotactic mutants in the first three days of soft agar migration, but then show a phenotype of enhanced migration in days 4-8. This enhanced migration was stable, suggesting that the *cheV1* mutants had acquired a suppressor mutation. These results thus suggest that *cheV1* mutants have strong soft agar migration defects, consistent with CheV1’s important role in forming chemoreceptor arrays (Abedrabbo *et al*., 2017).

After long soft agar incubation, cheV1 mutants accumulated SNPs in the HPG27_252 gene. The SNPs consisted of distinct insertions or deletions that resulted in a truncated open reading frame retaining from 23-60 of the full 179 amino acids, and lacking most of the predicted periplasmic domain. Complete deletion of HPG27_252 restored soft agar migration, supporting the hypothesis that removal of HPG27_252 allows enhanced soft agar migration of a *cheV1* mutant. The fact that a loss of function allows better soft agar migration fits well with the apparent high frequency of this occurrence.

It is not yet known why loss of HPG27_252 confers enhanced soft agar migration. HPG27_252 encodes a very remote homolog of TFF proteins that compose the cytoplasmic membrane alignment complex found in T4P and T2SS, PilO and GspM (McCallum *et al*., 2019; Naskar *et al*., 2021). Both PilO and GspM are cytoplasmic membrane proteins with a substantial periplasmic domain that adopts a ferredoxin-like fold. PilO and GspM form homodimers, as well as heterodimers with PilN/GspL, to create a cage-like ring structure. The exact function, however, remains unknown other than hypothesized ones to control TFF dynamics or align the cytoplasmic components with those at the outermembrane. The HPG27_252 gene is part of an operon with two other genes predicted to encode portions of the cytoplasmic membrane alignment complex, *pilN* (HPG27_251) and *pilM* (HPG27_250), and these genes are found in all *H. pylori* genomes. *H. pylori* does not have any reported T4P or T2SS, nor are any other genes for these processes near the locus identified here, suggesting that these TFF remote orthologs may be isolated and used for processes other than creation of TFF.

Recovery of soft agar migration is a powerful selection, and has led to the identification of new proteins involved in motility and chemotaxis (Partridge & Harshey, 2020). For example, Terry *et al*. used soft agar migration recovery in *H. pylori cheW* mutants to identify that *H. pylori* has a remote homolog of the CheZ chemotaxis phosphatase (Lertsethtakarn & Ottemann, 2010; Terry *et al*., 2006) and help to expand our understanding of the distribution of this protein (Liu *et al*., 2018). In some cases, the recovered strains are pseudo-revertants, meaning they occur outside the original targeted gene. Soft agar pseudo-revertants include mutations that affect flagellar reversals (Mohari *et al*., 2015) or ci-di-GMP levels (Nieto *et al*., 2019) as a way to override chemotaxis defects. We do not yet know how HPG27_252 allows soft agar migration. On one hand, it does not revert the extreme counterclockwise bias of Δ*cheV1* mutants. On the other hand, the protein does appear to interact directly with CheV1. Further work will be needed to untangle the role of HPG27_252, but this work has identified a new player in *H. pylori* chemotaxis and motility behavior.

## ACKNOWLEDGEMENTS

The authors would like to thank members of the Ottemann Lab, with special thanks to Skander Hathroubi and Kevin Johnson for their continued support throughout this project. We would also like to thank Andrew Kebbel for his guidance through the NGS data analysis process, and Thomas Hake for helping with certain computational statistical analyses in this study. The described project was supported by National Institutes of Health National Institute of Allergy and Infectious Disease (NIAID) grants number R01 AI116946 and R01 AI164682 (to K.M.O.). The funders had no role in study design, data collection and interpretation, or the decision to submit the work for publication.

## METHODS

### Bacterial culture conditions

*H. pylori* G27 strains (Censini *et al*., 1996) and *E. coli* strains used in this study are listed in Supplemental Table 2. *H. pylori* strains were grown on solid media consisting of Columbia horse blood agar with 5% v/v defibrinated horse blood (Hemostat Labs) with antibiotics at final concentrations of 50µg/ml cycloheximide, 10µg/ml vancomycin, 5µg/ml cefsulodin, 2.5units/ml polymyxin B, 5µg/ml trimethoprim, and 8µg/ml amphotericin B (CHBA). For mutant selection, chloramphenicol was used at 13.33ug/ml and kanamycin was used at 15ug/ml. The plates were incubated at 37°C in a microaerobic incubator with 10% v/v CO_2_, 5% v/v O_2_, and 85% v/v N_2_. Liquid cultures of *H. pylori* were grown in Brucella broth supplemented with 10% v/v heat-inactivated fetal bovine serum (FBS) supplemented with antibiotics mentioned above (BB10). *E. coli* strains were grown on LB plates with 100µg/ml ampicillin or 60µg/ml kanamycin antibiotics under normoxic conditions at 37°C. For long term storage, *H. pylori* was mixed with Brain Heart Infusion broth with 10% FBS, 1% (w/v) beta-cyclodextrin, 25% glycerol and 5% DMSO, and frozen at –80°C. To measure growth rates of Che+ suppressors, strains were grown in BB10 without antibiotics overnight.

### Soft-agar chemotaxis assays and isolation of Che^+^ suppressors

Soft-agar chemotaxis assays were done on Brucella broth plates supplemented with 0.35% Bacto Agar, 2.5% FBS, 50µg/ml cycloheximide, 10µg/ml vancomycin, 5µg/ml cefsulodin, 2.5units/ml polymyxin B, 5µg/ml trimethoprim, and 8µg/ml amphotericin B. Plates were allowed to set for 2-3 days at room temperature before inoculation to remove excess moisture. Plates were inoculated from cultures grown on CHBA plates. Plates were incubated at 37°C under microaerobic conditions and were imaged after 3, 5, and 7 days of incubation using the BioRad ChemiDoc XRS+ imaging system. For isolation of Che^+^ suppressor mutant colonies, samples were picked from the outer edge of the expanded *H. pylori* mutant colonies after 4-5 days of growth. These samples were restreaked on CHBA plates and colony purified.

### Next Generation Sequencing analysis

To purify genomic DNA for whole genome sequencing, *H. pylori* G27 wild-type, *cheV1* mutant, and four Che+ suppressors were grown on CHBA plates supplemented with chloramphenicol. Genomic DNA from relevant strains was purified using the Promega Wizard Genomic DNA Purification Kit using the protocol for Gram-negative bacteria provided by the manufacturer and sent for MiSeq sequencing at the UC Davis fee-for-service facility.

Whole genome sequencing (WGS) for the four suppressors through MiSeq resulted in 2×300 paired-end reads. To proceed further with next-generation sequencing (NGS) analysis, these reads were first checked for quality using FastQC v0.11.5 (https://www.bioinformatics.babraham.ac.uk/projects/fastqc/) to determine their quality scores and check for presence of Illumina sequencing adapters. BBDuk v37.25 (https://sourceforge.net/projects/bbmap/) was used to trim Illumina Truseq DNA adapters from the right end using a Kmer length of 15-27. BBDuk was also used to trim Low Quality bases from the 3’-end of the reads, removing bases lower than a quality score of Q20. Finally, reads less than 100bp were discarded. A quality score of Q20 was chosen as it corresponds to a 99% inferred base call accuracy by the sequencer. https://www.illumina.com/science/technology/next-generation-sequencing/plan-experiments/quality-scores.html

To determine variations through WGS data generated from the four Che^+^ suppressors, Geneious R11 (https://www.geneious.com) was used to map the quality controlled reads from each suppressor to the wild-type *H. pylori* G27 reference genome (Baltrus *et al*., 2008). Five iterations of mapping were used, with the Medium Sensitivity and High Sensitivity settings to confirm the results obtained were consistent and accurate. After mapping, variants were called using Geneious R11’s Find Variations/SNPs function, with a variant frequency of at least 75%, maximum variant P-value of 10^−6, and minimum strand-bias P-value set to 10^−5 when exceeding 65% bias.

### PCR primer design and DNA sequencing

PCR primers were designed using Geneious R11 (https://www.geneious.com). To confirm location of SNPs observed in whole genome sequencing, the region containing the gene HPG27_252 was amplified using primers 277,426F (5’-CCTTTAAGCGACGGGTGGTT) and 278,017R (5’-TTGCACCAACAGCCCTTCAT) in wild-type *H. pylori* G27, *H. pylori* G27 Δ*cheV1* mutant, Che+ Δ*cheV1*::*cat* Che+ Colonies 5, 6, 8, and 11. To confirm knockout of *cheV*1 after transformation in *H. pylori* G27 *cheV1* mutant, the surrounding region was amplified using primers HP_synDNA_277082F (5’-TGTGGTTTATGCACAACGCC) and HP_synDNA_279488R (5’-TGTAATGCCCGGACAGCTTT). Genomic DNA from these transformed cells was obtained following the protocol in Promega Wizard Genomic DNA Purification Kit (Cat# A1120).

### Mutant construction of ΔHPG27_252::kan

*H. pylori* HPG27_252 mutants were constructed by natural transformation with plasmid pHPG27_252::*aphA3* DNA (Supplemental Table 2), followed by selection for kanamycin resistance. pHPG27_252::aphA3 was constructed synthetically (Genewiz) to knockout HPG27_252 by replacement with *aphA*3 leaving a 24bp and 19bp of the original gene remaining at the 5’-end and 3’-end respectively. Successful mutants were confirmed by PCR using primers flanking HPG27_252 (277320F (GCAAAGCGTGGTGGTTAGTG) and 277980R (GCAAAGCGTGGTGGTTAGTG) and sent for Sanger sequencing.

### Bacterial two hybrid analysis

Full-length of HPG27_252 gene was amplified using the *H. pylori* strain G27 genomic DNA as template with primer pairs P97/P98, in introduce into pUT18 and P105/P106, to introduce into pUT18C. The sequences are as follows, with engineered restriction sites shown in lowercase:

P97 (CCCaagcttATGAAGCCATTGCATTTTTCACACTTAGAC)
P98 CGggatccCTATTGGAGATCAATACTCACGCTAAATTGCA
P105 GCtctagaATGAAGCCATTGCATTTTTCACACTTAGAC
P106 GGggtaccCTATTGGAGATCAATACTCACGCTAAATTGCA

The amplicons were introduced into BACTH plasmids pUT18 and pUT18C withT4 DNA ligase after being digested with cognate restriction enzymes. The resulting plasmids were transformed into *E. coli* DH10B using LB plates with 100 μg/mL ampicillin. The colonies were colony purified, and then the correct plasmids were confirmed by amplifying HPG27_252 using primers P97 and P98. These plasmids are called pUT18-HPG27_252 and pUT18C-HPG27_252. Plasmid pKNT25-cheV1 was purified from strains previously described (Abedrabbo *et al*., 2017). *E. coli* BTH101 (Supplemental Table 2) was transformed with either pUT18-HPG27_252 and pKNT25-cheV1, or pUT18C-HPG27_252 and pKNT25-cheV1. Positive controls were plasmids pKT25-zip and pUT18c-zip, expressing leucine zipper motifs and negative controls were pKT25 and pUT18C (Supplemental Table 2). Co-transformed cells were plated on LB plates with 100μg/mL ampicillin and 30 μg/mL kanamycin. Resulting colonies were then cultured overnight in liquid, and then 10 µl was dropped on to LB plates with 100 μg/mL ampicillin, 30 μg/mL kanamycin, 40 μg/mL X-gal, and 0.5 mM IPTG and incubated at 30 °C. Representative photos were taken after culturing for 40 h.

### Bioinformatics Analysis of HPG27_252

Gene and protein sequences were obtained from the UCSC Microbial genome browsers. Operon structure was obtained from Sharma *et al*. (Sharma *et al*., 2010), as visualized on the UCSC 26695 genome browser (Chan *et al*., 2012). Transmembrane domain and topology predictions were done with Deep TMHMM (Hallgren *et al*., 2022). Remote homology detection was done using HHPRED at the MPI Bioinformatics Toolkit (Zimmermann *et al*., 2018).

### Live cell tracking

For live cell tracking assays, liquid cultures of *H. pylori* were grown overnight in BB10 and diluted to OD600s between 0.2 and 0.3 in fresh BB10 before and filmed under a microscope the next day. Movies were captured using a Nikon Eclipse E600 microscope connected to a Hamamatsu 1394 ORCA-285S camera using a 20X phase-contrast objective and μManager (Edelstein *et al*., 2014). One minute of video was captured with an approximate frame rate of 13 frames per second. These movies were then analyzed in 3 second intervals using ImageJ v1.53 (Schneider *et al*., 2012) for flagellar reversals/direction changes which were characterized as a sharp change of cell direction. A total of 100 cells were quantified for each strain used for this assay.

### Growth analysis

Liquid overnight cultures were grown of wild-type *H. pylori* G27, *cheV1* mutant, and Che+ colony 5 in BB10. After overnight growth, the cultures were diluted to an OD600 of 0.250 using fresh BB10, and OD600 values were measured every 3 hours for 12 hours.

## Supplemental

**Supplemental Fig. 1:**
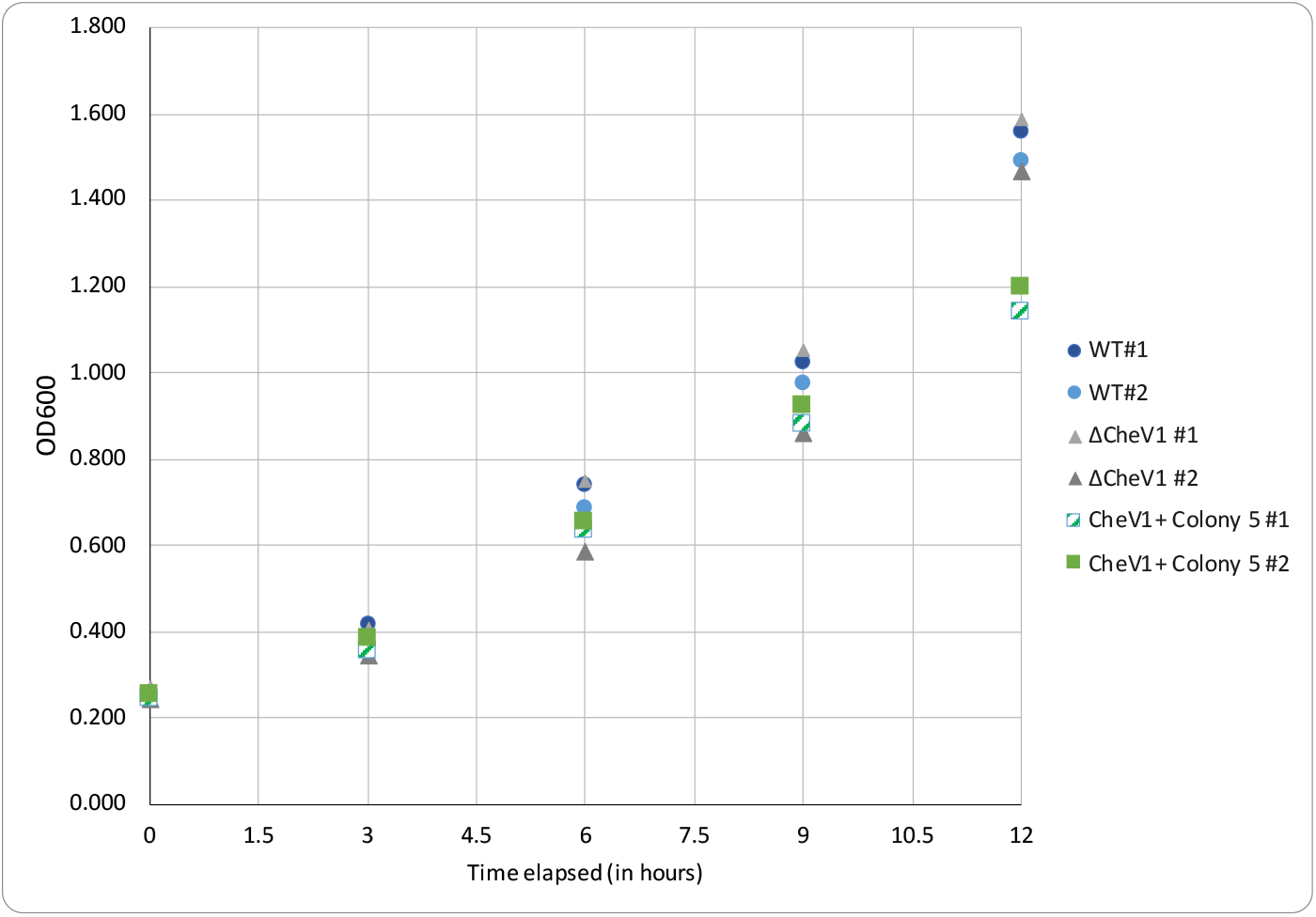
Growth measurements. Growth curve measurements of H. pylori G27 WT, ΔcheV1, and ΔcheV1 Che+ colony 5. Each strain was analyzed in biological duplicates.

**Supplemental Fig. 2.**
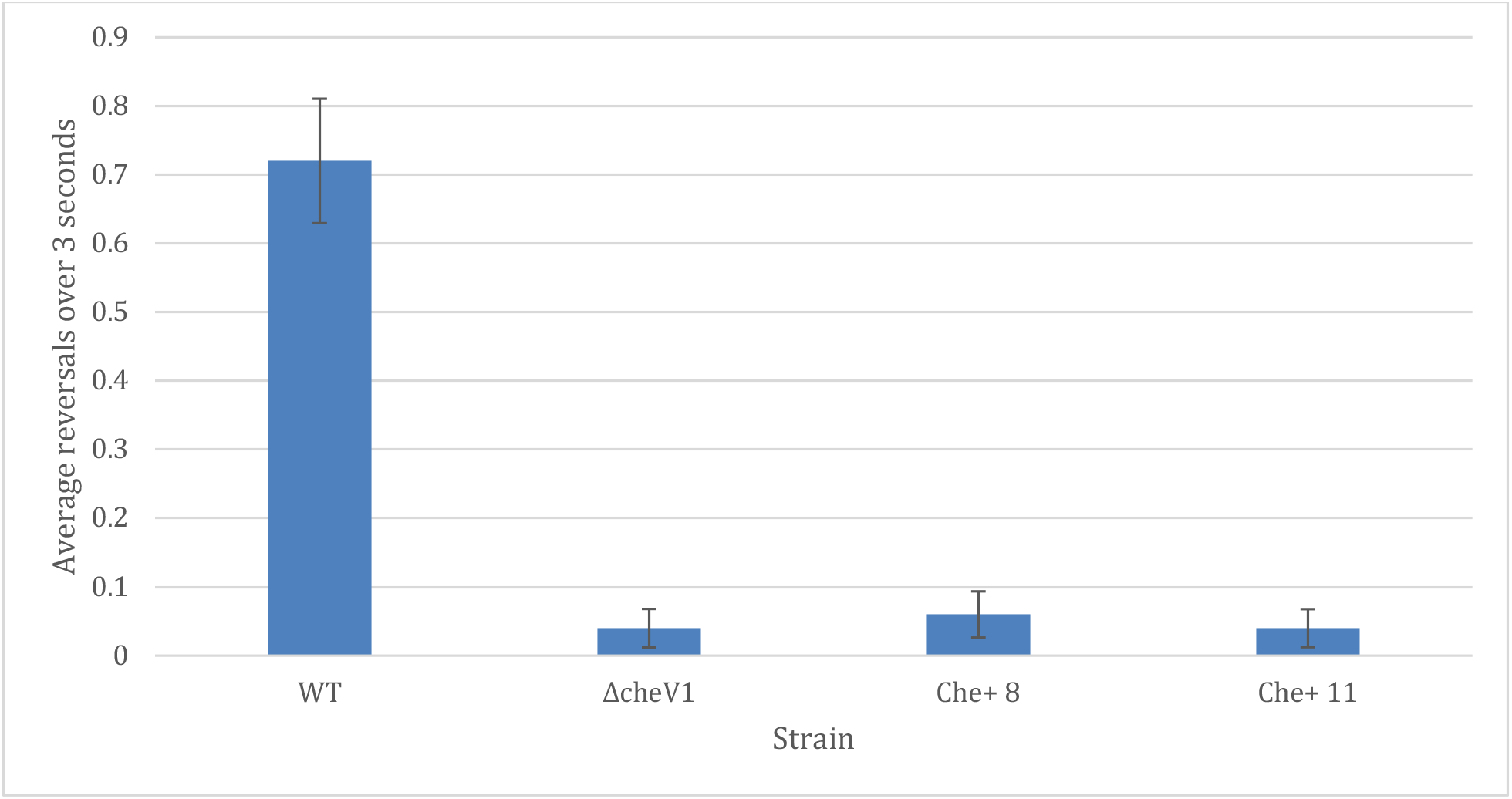
reversal phenotype. Swimming reversal frequency of *H. pylori* WT, ΔcheV1 and two Che+ colonies. Average reversals was determined by filming liquid cell cultures, following the trajectory of an individual cell for at least 3 seconds, and manually identifying reversals. These were enumerated for 50 cells for each strain over 2 biological replicates. Error bars represent the standard error of the mean.

**Supplemental Table 1:**
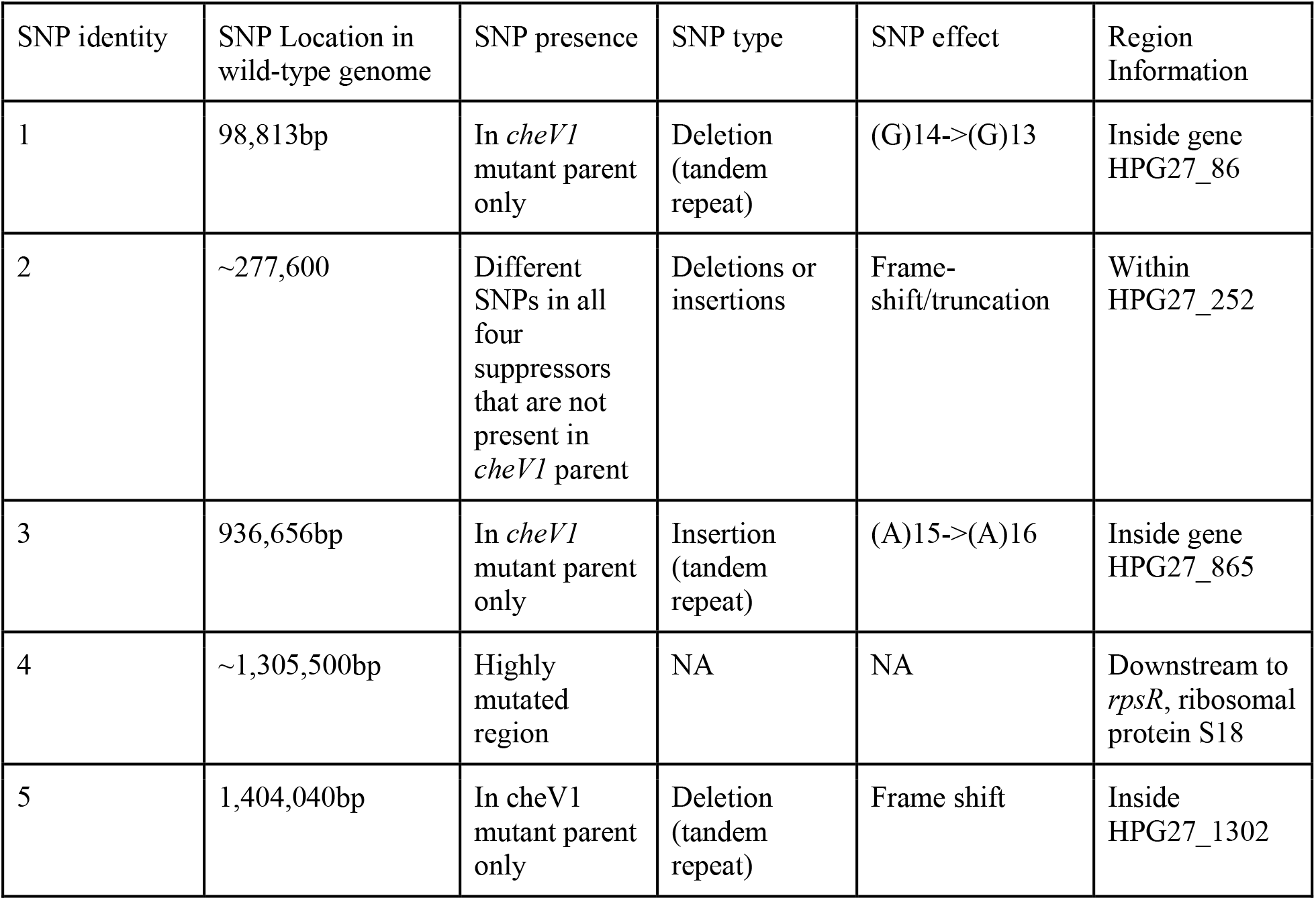
SNPs. SNPs identified in the sequencing of *H. pylori* G27 *cheV1* and its four Che^+^ suppressors. The region for SNP #1 was Sanger sequenced to confirm the existence of this SNP. Based on the Sanger sequencing data, it was determined that this SNP was an artifact of sequencing due to the poly-G region and was therefore discarded from further analysis. SNP #2 was then considered the most promising candidate as the SNPs varied on each single suppressor colony. Sanger sequencing from this region confirmed this mutation in all suppressors.

| SNP identity | SNP Location in wild-type genome | SNP presence | SNP type | SNP effect | Region Information |
| --- | --- | --- | --- | --- | --- |
| 1 | 98,813bp | In *cheV1* mutant parent only | Deletion (tandem repeat) | (G)14->(G)13 | Inside gene HPG27_86 |
| 2 | ~277,600 | Different SNPs in all four suppressors that are not present in *cheV1* parent | Deletions or insertions | Frame-shift/truncation | Within HPG27_252 |
| 3 | 936,656bp | In *cheV1* mutant parent only | Insertion (tandem repeat) | (A)15->(A)16 | Inside gene HPG27_865 |
| 4 | ~1,305,500bp | Highly mutated region | NA | NA | Downstream to *rpsR*, ribosomal protein S18 |
| 5 | 1,404,040bp | In *cheV1* mutant parent only | Deletion (tandem repeat) | Frame shift | Inside HPG27_1302 |

**Supplemental Table 2.** Strains and plasmids used in this study. *H. pylori* and *E. coli* strains and plasmid used in this study

| Organism | Strain | Relevant phenotype | KO Lab Number | Source |
| --- | --- | --- | --- | --- |
| *H. pylori* | G27 Wild-type | Motile, chemotactic (Che+) | KO379 | Nina Salama/(Censini *et al.*, 1996) |
| | G27 $\Delta che1::cat$ | Motile, non-chemotactic (Che-) | KO1277 | (Lertsethtakarn *et al.*, 2015) |
| | G27 $\Delta cheV1::cat$ Che+ Colony 5 | Che+ | KO1458 | This study |
| | G27 $\Delta cheV1::cat$ Che+ Colony 6 | Che+ | KO1459 | This study |
| | G27 $\Delta cheV1::cat$ Che+ Colony 8 | Che+ | KO1461 | This study |
| | G27 $\Delta cheV1::cat$ Che+ Colony 11 | Che+ | KO1464 | This study |
| | G27 $\Delta HPG27\_252::aphA3$ | Che+ | KO1700 | This study |
| *E. coli* | DH10B Wild-type | Cloning strain | KO14 |  |
|  | BTH101 | Bacterial two hybrid strain |  |  |
| Plasmids | DH10B pHPG27_252::aphA3 | Deletion plasmid for HPG27_252 | KO1699 | This study |
|  | pUT18 |  | KO1346 | (Karimova *et al.*, 1998) |
|  | pUT18C |  | KO1347 | (Karimova *et al.*, 1998) |
|  | pKNT25-cheV1 | BACTH plasmid with T25 fused to CheV1 | KO1431 | (Abedrabbo *et al.*, 2017) |
|  | pUT18-HPG27_252 | BACTH plasmid with T18 fused to the C-terminus of HPG27_252 |  | This study |
|  | pUT18C-HPG27_252 | BACTH plasmid with T18 fused to the N-terminus of HPG27_252 |  | This study |
|  | pKT25-zip |  | KO1348 | (Karimova *et al.*, 1998) |
|  | pUT18c-zip |  | KO1349 | (Karimova *et al.*, 1998) |

